# Landscape heterogeneity buffers biodiversity of meta-food-webs under global change through rescue and drainage effects

**DOI:** 10.1101/2020.06.03.131425

**Authors:** Remo Ryser, Myriam R. Hirt, Johanna Häussler, Dominique Gravel, Ulrich Brose

## Abstract

The impacts of habitat fragmentation and eutrophication on biodiversity have been studied in different scientific realms. Metacommunity research^1–5^ has shown that reduction in landscape connectivity may cause biodiversity loss in fragmentated landscapes. Food-web research addressed how eutrophication increases biomass accumulations at high trophic levels causing the breakdown of local biodiversity ^6–9^. However, there is very limited understanding of their cumulative impacts as they could amplify or cancel each other. Here, we show with simulations of meta-food-webs that landscape heterogeneity provides a buffering capacity against increasing nutrient eutrophication. An interaction between eutrophication and landscape homogenization precipitates the decline of biodiversity. We attribute our results to two complementary mechanisms related to source and sink dynamics. First, the “rescue effect” maintains local biodiversity by rapid recolonization after a local crash in population densities. Second, the “drainage effect” allows a more uniform spreading of biomass across the landscape, reducing overall interaction strengths and therefore stabilizing dynamics. In complex food webs on large spatial networks of habitat patches, these effects yield systematically higher biodiversity in heterogeneous than in homogeneous landscapes. Our meta-food-web approach reveals a strong interaction between habitat fragmentation and eutrophication and provides a mechanistic explanation of how landscape heterogeneity promotes biodiversity.

---

Increasing human demands for production of goods in natural landscapes have caused habitat fragmentation and homogenisation, eutrophication and increasing land-use intensity. This resulted in an erosion of biodiversity and associated ecosystem services at global scales. Habitat fragmentation describes how production areas dissect continuous natural landscapes into habitat patches embedded in a landscape matrix whose hostility for the species increases with land-use intensity. Increasing nutrient inputs from agricultural practices yield biomass accumulations at higher trophic levels, eroding biodiversity by increased species’ interaction strengths^6,9^. Despite growing evidence on the importance of these global change factors, we still do not understand how their interaction drives biodiversity changes. While fragmentation and eutrophication are often studied in isolation, complex feedback loops in multi-trophic food webs can generate non-linearities in the response of biodiversity, which is rendering our knowledge of the interactive effects of these stressors in natural landscapes fraught with uncertainty. The high-dimensional interplay between spatial and trophic processes prevents experimental studies on such complex interactions. Simulations of spatial food web dynamics are therefore needed to reveal the mechanisms underlying how these global change stressors interact.

One key challenge is the integration of spatial processes connecting local populations across habitat patches into metapopulations and interaction processes connecting local species into complex food webs (Fig. 1). Traditionally, independent and mostly separated research areas have addressed these two types of ecological networks. First, metacommunity theory describes how dispersing individuals connect local populations across complex spatial networks of habitat patches^10^. Depending on their size and quality, patches can comprise large source populations that yield a net dispersal flux of individuals to small sink populations^1,4^ (Fig. 1a). These source-sink dynamics^11^ can facilitate persistence of small populations by rescue effects^12^, which is undermined by increasing fragmentation or land-use intensity that prevent successful dispersal. Second, food-web theory addresses how biomass fluxes (i.e. energy and matter) between species drive population dynamics (Fig. 1b). Weak biomass fluxes can cause consumer extinction due to energy limitations while strong biomass fluxes can result in top-heavy consumer-resource biomass pyramids with unstable dynamics^6,9^. Eutrophication in particular increases all biomass fluxes and thus undermines biodiversity of local food webs^7^. Although both research areas documented strongly negative effects of either fragmentation or eutrophication on biodiversity, the interplay of these stressors in complex natural communities has remained virtually untapped.

**Fig. 1.**
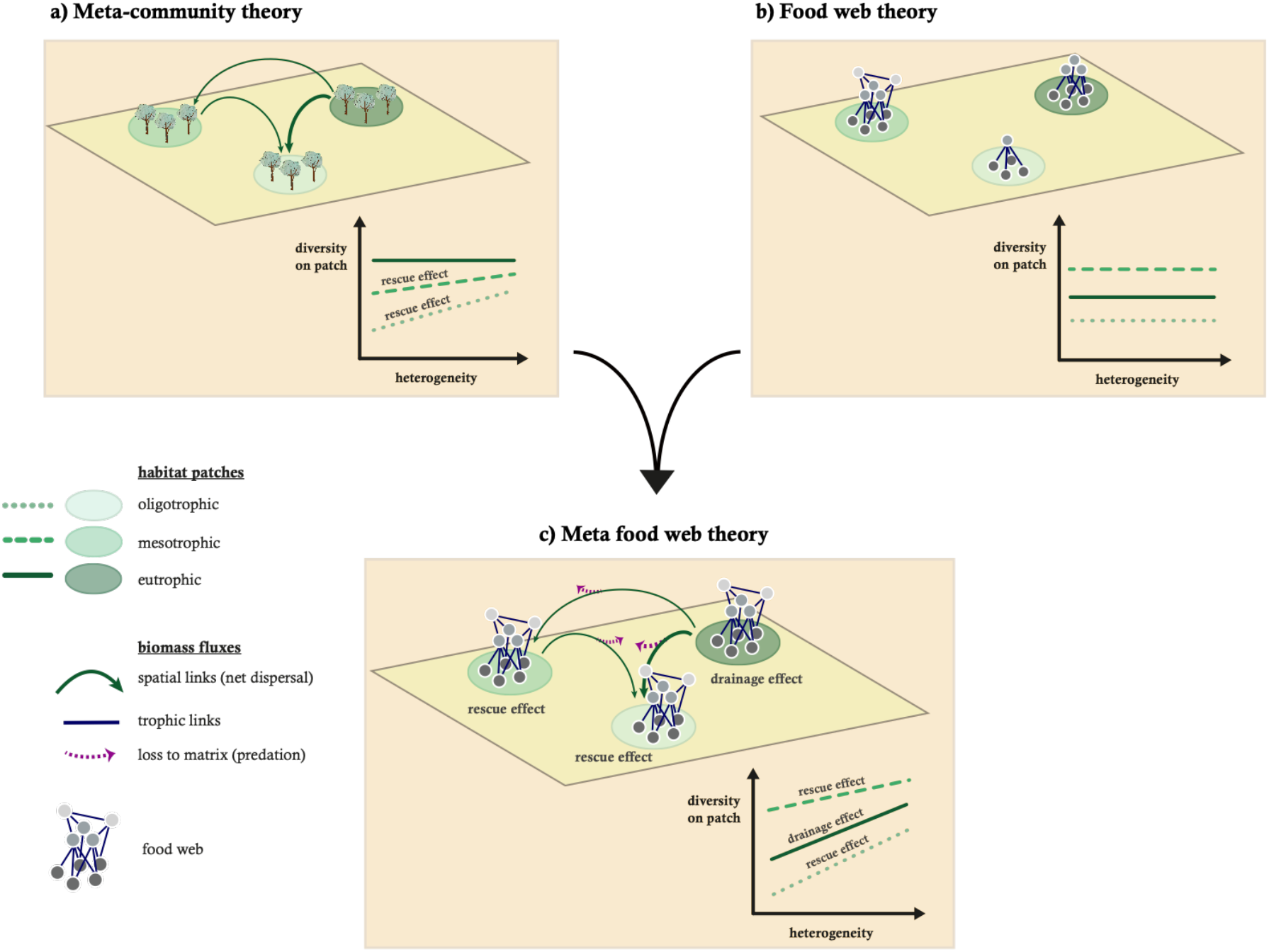
Conceptual figure illustrating the synthesis of metacommunity theory and food-web theory into meta-food-web theory. Panel a) illustrates metacommunity dynamics with net dispersal from larger (nutrient richer patches) to smaller populations (nutrient poorer patches) and the associated rescue effect on local diversity. Panel b) illustrates local food-web dynamics on patches with different nutrient richness and the effect of the paradox of enrichment on local diversity. Panel c) illustrates the synthesis of metacommunity and food web dynamics and the interaction of respective key effects and their consequence for biodiversity.

So far, studies synthesizing spatial and trophic processes have been limited to small species motifs such as food chains^13,14^. They showed that dispersal can synchronize population dynamics, which reduces biodiversity by correlated local extinctions^15,16^. However, consumer dispersal can also induce compensatory dynamics^17^ and dampen oscillations^18^, which prevents extinctions. Moreover, dispersal may increase resilience of complex food webs by reducing strong interspecific interactions^19^ depending on the trophic level that is dispersing^18^. The relative strength of these potentially counteracting positive and negative effects of dispersal on population persistence depends on the trophic interaction structure^14^. While these studies have demonstrated interactions between spatial and trophic processes in small modules, the study of impacts on biodiversity in large spatial networks with many species has remained in its infancy. Traits of organisms play an important role in both spatial and trophic processes. In metacommunities, body mass and movement mode determine which patches compose species-specific spatial networks^20^. Similarly, the propagation of energy fluxes through food webs is driven by species’ interaction strengths that depend strongly on body masses^8^. Although metapopulation and food-web theories have been developed mostly independently, they have identified the same important drivers (i.e. body mass), and the same currencies (i.e. biomass fluxes). To date, a trophic metacommunity framework incorporating spatial use properties is still lacking^21^. Also, as spatial and trophic processes in real landscapes are coupled (Fig. 1c), a mechanistic understanding of global change effects on ecosystems will benefit from an integrated approach. We address this challenge by synthesizing metapopulation and food-web models that use allometric scaling relationships of spatial and trophic processes as a unifying principle into a meta-food-web model. We identify key mechanisms complementary to the rescue effect in landscapes under eutrophication and isolation (Fig. 1).

We use a bioenergetic model to analyse population dynamics across a gradient of complexity from simple (tri-trophic food chain on a single patch) to complex systems (40-species food web on 50 habitat patches). This model employs body masses as the unifying trait that determines not only trophic links and interaction strengths of the food webs but also the dispersal ranges. Dispersal rates depend on local net growth rates, summarizing resource availability, competition and predator pressure arising from local trophic dynamics^22^.

Firstly, on a single patch, low nutrient supply for a tri-trophic food chain causes predator starvation (Fig. 2a, extinction, left side). Increasing nutrient supply first promotes predator equilibrium biomass densities (Fig. 2a, survival, equilibrium) and therefore top-heavy biomass pyramids causing biomass oscillations (Fig. 2a, survival, oscillation), which paradoxically eventually yield predator extinction (Fig. 2a, extinction, right side). Such extinctions due to unstable oscillations under eutrophication have first been described as the “paradox of enrichment”^6^. Subsequently, they were generalized to systems with an increased energy flux to the predator relative to its loss rate^9,23^. Turning around this “principle of energy flux”, however, also suggests that an additional drainage effect arises from energy transfer from large populations (sources) to small populations (sinks), preventing unstable dynamics in top-heavy systems. Consistent with this hypothesis, we find that increasing emigration rates that drain biomass out of a eutrophic location can prevent predator extinction by reducing oscillations (Fig. 2b). Spatial fluxes tend to increase with dispersal rates and the underlying variability in the landscape. This demonstrates the drainage effect as a mechanism by which spatial processes can stabilize trophic population dynamics in heterogeneous landscapes.

**Fig. 2:**
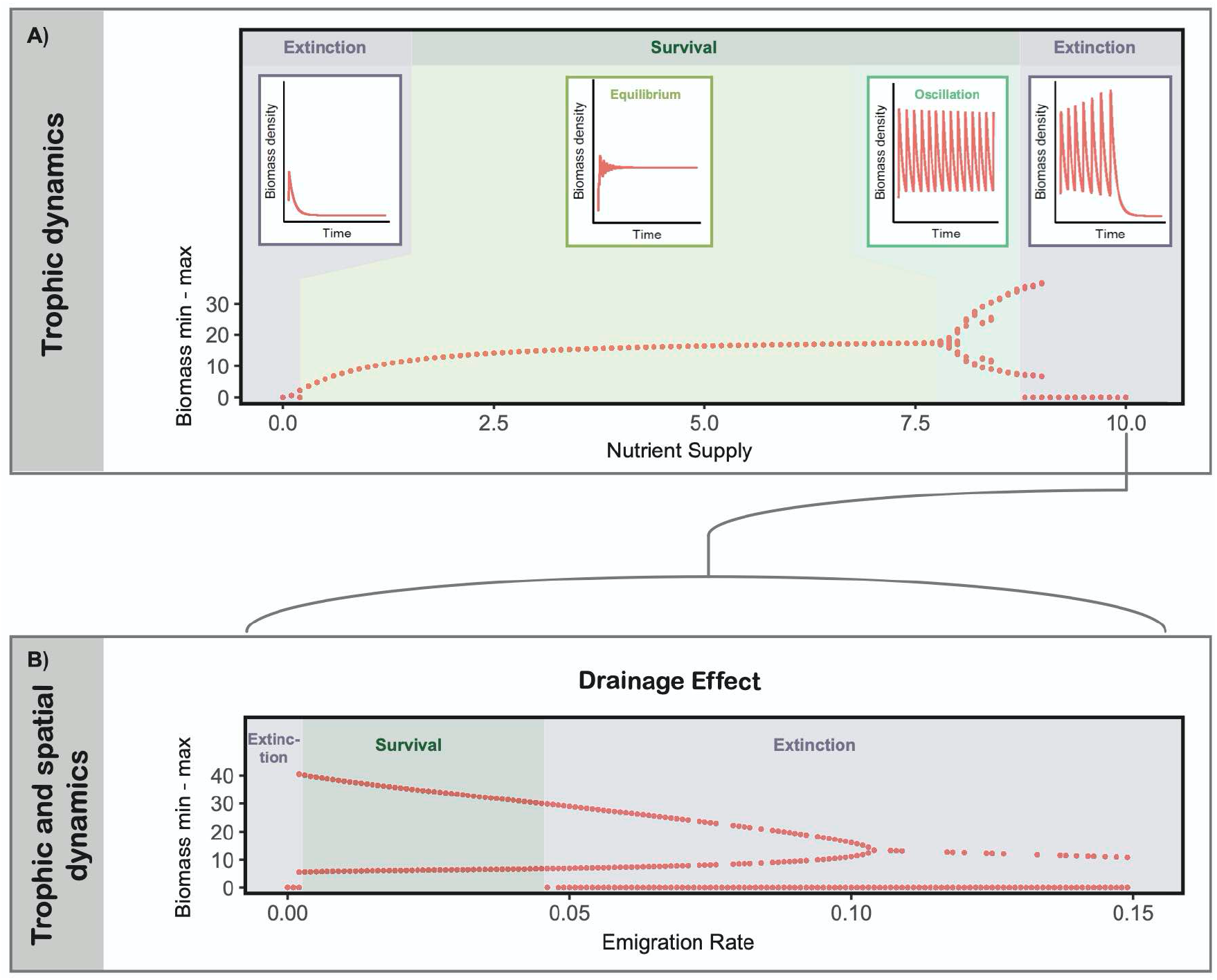
Top predator dynamics of a tri-tropic food chain on a single patch. A) Exemplary time series of biomass densities of the predator at different nutrient supply concentrations (boxes; from left to right: 0.1 (oligotrophic); 3 (mesotrophic); 8.5 and 10 (eutrophic)) corresponding to points in the bifurcation diagram showing maximum and minimum biomass density (y-axis) across a gradient of nutrient supply concentrations (x-axis). B) Bifurcation diagram showing maximum and minimum biomass density (y-axis) when enabling emigration across a gradient of emigration rates (x-axis) with a nutrient supply concentration of 10, which corresponds to the last point in panel A).

Subsequently, we studied this drainage effect in systems of two connected habitats across gradients of landscape hostility and habitat heterogeneity (represented by the difference in nutrient supply concentration of the two locations). Landscape hostility summarizes all factors that drive the loss of biomass during dispersal including higher metabolic costs and increased mortality rates by predation pressure in the unsuitable landscape matrix. Dispersal synchronizes unstable dynamics, causing predator extinction (Fig. 3, lower left corner), in simulations without heterogeneity and without hostility. Increasing landscape hostility yields drainage of biomass during dispersal, facilitates predator persistence and then also reduces oscillations (Fig. 3, along the hostility axis). At very high levels of landscape hostility, however, extreme death rates during dispersal cause predator extinction. Similarly, increasing patch heterogeneity also enables predator persistence and decreases oscillations (Fig. 3, along the heterogeneity axis). The drainage effect offers general mechanistic explanations for these emergent patterns despite of some slightly more complex patterns in population oscillations (e.g. some combinations of landscape hostility and patch heterogeneity yield weak spatial links between patches and desynchronization of biomass oscillation frequencies, see Supplement Fig. S2 for details). For eutrophic patches, increased dispersal losses by landscape hostility or the coupling with an oligotrophic patch (patch heterogeneity) both increase the biomass drainage through increased net migration. For oligotrophic patches, however, there are differences between effects of landscape hostility and patch heterogeneity. Drainage by landscape hostility supresses small populations even more, whereas patch heterogeneity causes a gain in biomass via dispersal that supports predator populations via rescue effects (see Supplement Fig. S1). Patch heterogeneity thus creates dispersal fluxes in biomass that are responsible for not only the well-known rescue effects^12^ supporting small populations on oligotrophic sink patches by net-immigration but also the drainage effects sustaining large populations on eutrophic patches.

**Fig. 3:**
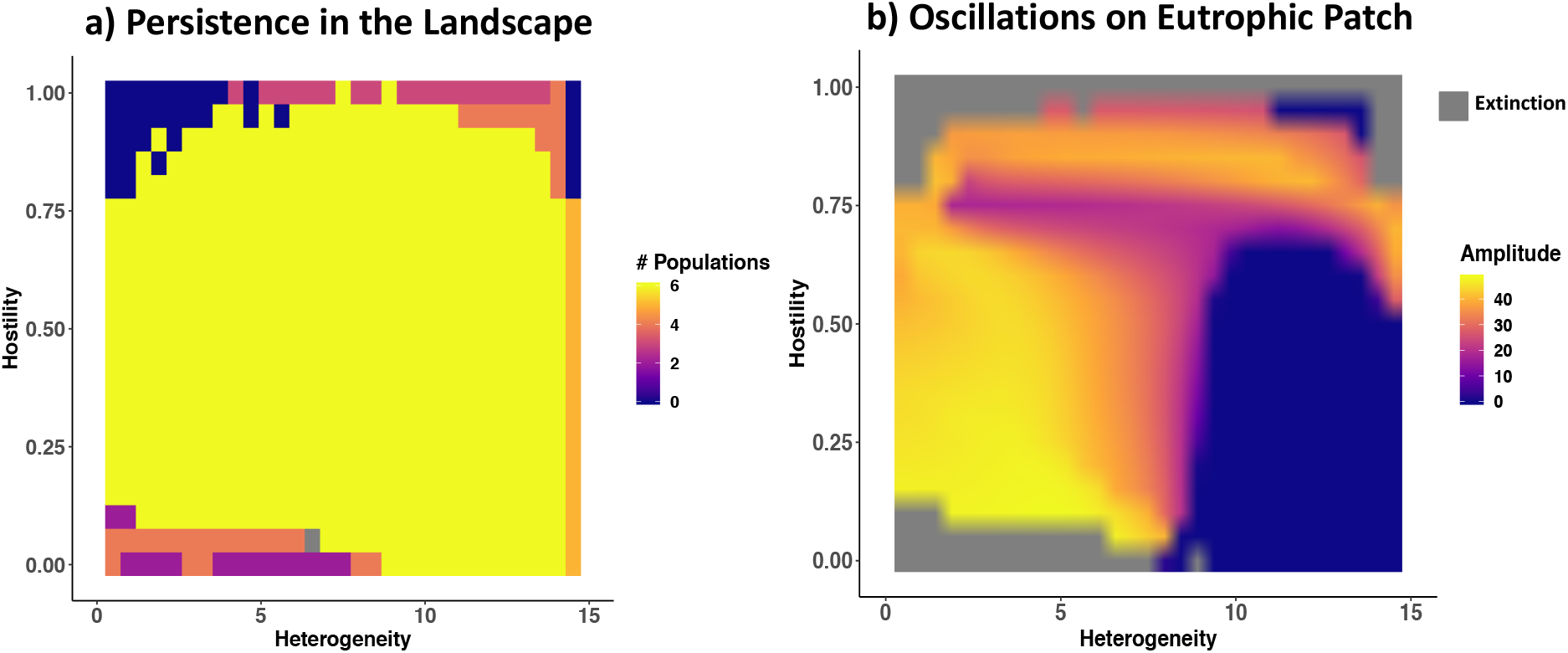
Top predator dynamics of a tri-tropic food chain on two coupled patches. a) Heat map showing the number of persisting populations (colour coded; plant, herbivore and predator on 2 patches; maximum of 6) in the landscape across gradients of landscape heterogeneity (x-axis; difference in nutrient supply concentration across the two patches; on the left: two eutrophic patches, on the right: an eutrophic and an oligotrophic patch) and matrix hostility (y-axis). b) Heat map showing the amplitude of biomass density oscillations of the predator (z-axis; colour coded) in the (always) eutrophic patch across gradients of landscape heterogeneity (x-axis; difference in nutrient supply concentration between the two patches) and matrix hostility (y-axis). Amplitudes of 0 (blue) stand for an equilibrium state of the predator. Grey areas are where the predator went extinct.

To generalize the mechanistic understanding of drainage effects from food chains, we simulated the dynamics of a complex food web consisting of 10 plants and 30 animals on different complex landscapes containing 50 habitat patches (Fig. 4). We simulated homogeneous landscapes, where all patches have the same nutrient supply concentration. These simulations were replicated across a gradient of nutrient supply concentrations ranging from 10^−0.8^ (oligotrophic) to 10^2^ (eutrophic). We also simulated three types of heterogeneous landscapes with landscape averages being oligotrophic, mesotrophic or eutrophic (Fig. 4). Nutrient supply concentration for each patch of heterogenous landscapes is assigned randomly from the same gradient as in the homogeneous scenario, but with a higher sampling density in the lower or higher nutrient supply values for oligotrophic and eutrophic heterogeneous landscapes, respectively, and uniform sampling for the mesotrophic heterogeneous landscapes. In line with our results from the food chain simulations, we found that local species richness in homogeneous landscapes is lowest on oligotrophic patches due to energy limitation. Higher nutrient supply first increases species richness on mesotrophic patches before decreasing it again on eutrophic patches (Fig. 4, purple). Species richness is highest in mesotrophic heterogeneous landscapes because oligotrophic patches profit from the rescue effect and eutrophic patches profit from the drainage effect (Fig. 4, orange). If there are only a few oligotrophic patches in a eutrophic heterogeneous landscape, rescue and drainage effects still increase local diversity, although the recue effect is weaker (Fig. 4, blue). Similarly, a few eutrophic patches in an oligotrophic landscape foster local diversity through rescue and drainage effects (Fig. 4, green). Thus, rescue effects and drainage effects also apply to complex food webs in complex landscapes. This shows that the interaction of strong and weak spatial and trophic biomass fluxes increases stability and species richness in metacommunities.

**Fig. 4.**
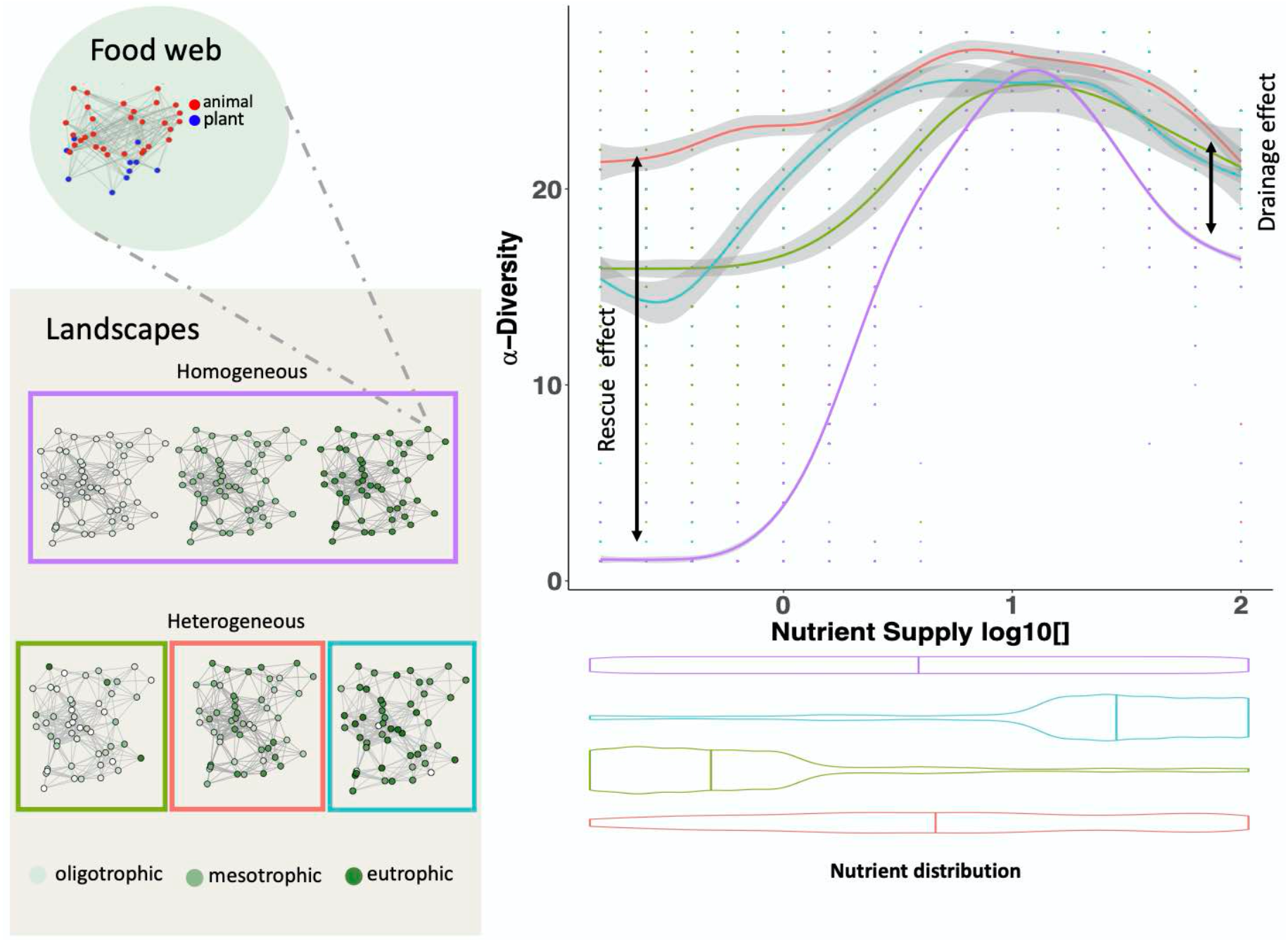
Landscape heterogeneity drives biodiversity in complex meta-food-webs. Local diversity (y-axis) across a gradient of patch nutrient supply concentration in homogeneous (purple) and heterogeneous (green, orange, blue) landscapes. Violin plots below the x-axis show nutrient distributions within the landscape for each scenario. The meta-food-web consists of a complex food web of 10 plants and 30 animals and large homogeneous and heterogeneous landscapes with 50 habitat patches with different patch nutrient supply concentrations (nutrient supply concentrations on habitat patches are colour coded). Edges indicate dispersal links for an exemplary species with a dispersal range of 0.3. Lines are a smooth fit from a GAM model with 95% confidence intervals in ggplot2 and points represent the data.

Spatial processes in heterogenous landscapes stabilise local food-web dynamics and translate into higher diversity. This stresses the importance of addressing global change drivers in a meta-food-web framework. Various mechanisms are involved, all related to source-sink dynamics where energy moves from high biomass locations to low biomass locations. We have found that the well-known rescue effect allows persistence on oligotrophic patches, while the novel drainage effect buffers eutrophic patches. Complex interactions among these phenomena may further promote diversity. For instance, nutrient spillover from a eutrophic to a neighbouring oligotrophic location may promote local productivity and increase food-chain length^24^. Such spatial nutrient diffusion can destabilize simple food chains and decrease spatial heterogeneity in a meta-ecosystem model^18^ and thus cross-ecosystem nutrient fluxes can change community composition^25^. These meta-ecosystem approaches have synthesized nutrient fluxes with simple trophic modules, and our meta-food-web approach provides a flexible tool to scale-up these findings to the levels of landscape and food-web complexity that characterize natural ecosystems.

In real landscapes, which suffer more and more from fragmentation, land-use intensification and eutrophication due to human activities, managing connectivity and heterogeneity is an important aspect of biodiversity conservation and restoration. Traditionally, increasing landscape hostility due to higher dispersal mortality or increased distances between habitat fragments have been perceived as threats to the biodiversity of habitat patches as they reduce rescue effects^12^. Hence, wildlife bridges across highways and other corridors to increase connectivity between habitat patches have been propagated as important tools to remedy the consequences of land-use intensification as the reduced hostility may benefit small sink populations by rescue effects and thus lower extinction risks^26^. Our results, however, indicate that the consequences of increasing habitat connectivity are highly context-dependent. We found that higher connectivity between large populations can undermine biodiversity by decreasing the drainage effect, whereas connecting large and small populations is generally beneficial for both. Thus, in managing landscape connectivity, connections between eutrophic and oligotrophic habitats or among oligotrophic habitats should be enhanced to reduce the hostility effect. However, connections among eutrophic habitats should only be established with caution, as a reduced hostility effect results in less drainage effect and thus has the potential do destabilize both populations.

Broader implications for ecosystem services can arise as two habitat patches that suffer from eutrophication may lose predatory pest control agents if they are well connected to each other but may maintain pest control when coupled with less intensive or natural habitats. Thus, the management of connectivity and heterogeneity in landscapes suffering from fragmentation and eutrophication may benefit from fostering rescue and drainage effects to maintain biodiversity and ecosystem services. Our meta-food-web approach has revealed interactions between spatial and trophic dynamics beyond the rescue effect that provide a mechanistic explanation of how landscape heterogeneity enhances biodiversity, which facilitates new strategies for active landscape management to foster natural biodiversity and ecosystem services.

## Author contributions

R.R., U.B. and M.R.H. developed the idea, R.R. built the model, did the analyses and wrote the first draft of the manuscript. M.R.H. designed the figures. All authors contributed to the interpretation and the final version of this manuscript.

## Acknowledgements

We gratefully acknowledge the support of the German Centre for Integrative Biodiversity Research (iDiv) Halle-Jena-Leipzig funded by the German Research Foundation (FZT 118) and funding by the German Research Foundation (DFG) in the framework of the research units FOR 1748 (BR 2315/16-2) and FOR 2716 (BR 2315/21-1).

## Materials and Correspondence

Requests should be addressed to U.B.

## Methods summary

### Model

We model a tritrophic food chain of one plant, one herbivore and one predator population on one or two habitat patches and complex meta-food-webs consisting of 10 plants and 30 animals in different landscapes containing 50 patches. The feeding dynamics are constant over all patches and are determined by the allometric food-web model by Schneider et al. 2016^27^. We integrate dispersal as species-specific biomass flux between habitat patches according to Ryser et al. 2019^28^. With the use of a dynamic bioenergetic model we formulate feeding and dispersal dynamics in terms of ordinary differential equations. The rate of change in biomass densities of a species are the sum of its biomass loss by metabolism, being preyed upon and emigration and its biomass gain by feeding and immigration. For detailed equations see Ryser et al. 2019^28^ and for model parameters see the supplement (TS1).

### Local food-web dynamics

Following the allometric food-web model by Schneider et al. 2016^27^ each species is fully characterised by its average adult body mass. For the complex food web log_10_ body masses were randomly drawn from a uniform distribution from 0 to 3 for plants and from 2 to 6 for animals. For the food chain the plant body mass was set to 10^2^, the herbivore body mass to 10^4^ and the predator body mass to 10^6^. We set mass ratios of the herbivore to the plant and the predator to the herbivore to the optimum of 100, thus the respective resource being a one-hundredth of its consumer’s body mass. Trophic dynamical parameters, such as metabolic rates and feeding rates, scale with body masses of model species. Also, we assume a type II functional response. Capture rates were reduced to 5% to achieve viable food chains and food webs with no interference competition.

### Nutrient model

We have an underlying nutrient model with one nutrient that is driving the nutrient uptake and therefore the growth rate of the plant population^8,27^. The nutrient model consists of one nutrient, a nutrient turnover rate of 0.25 and a nutrient supply concentration. The nutrient supply concentration was varied to get eutrophic and oligotrophic patches (see Setup).

### Spatial dynamics

We model dispersal between local communities as a dynamic process of emigration and immigration, assuming dispersal to occur at the same timescale as the local population dynamics^29^. Thus, biomass flows change dynamically between local populations and the dispersal dynamics directly influence local population dynamics and vice versa^22^.

Dispersal rates of animals are modelled with an adaptive emigration rate depending on the net growth rate on the given patch. Dispersal ranges depend on the body masses of our model species with larger species having a higher dispersal range. We model a hostile matrix between habitat patches that does not allow feeding interactions to occur during dispersal. Depending on the scenario, we define a landscape with one or two patches. In cases with two patches, their locations are spatially explicit and were chosen in a way that the distances between reflect the dispersal loss of the predator across the matrix hostility gradient.

### Emigration and immigration

Based on empirical observations^30^ and previous theoretical frameworks^13,20,31^, we assume that the maximum dispersal distance of animal species increases with their body mass. For simplicity, we do not let the plants disperse, as they don’t move themselves and the dispersal of plant propagules strongly depends on their dispersal strategy. We model emigration rates as a function of each species’ per capita net growth rate, which is summarising local conditions such as resource availability, predation pressure, and inter-and intraspecific competition^22^. Dispersal losses scale linearly with the distance between two patches and are 100% in scenarios with only one patch or when the distance between the two patches surpasses the dispersal range of an animal. Even though we model dispersal losses according to dispersal distances, this loss term could also represent any other sort of dispersal loss. For numerical reasons, we did not allow dispersal flows smaller than 10^−10^.

### Numerical simulations

We initialised each local population with a biomass density randomly sampled from a uniform probability density within the interval (0,10). Starting from these random initial conditions, we numerically simulated food web and dispersal dynamics over 100,000 time steps by integrating the system of differential equations implemented in C++ using procedures of the SUNDIALS CVODE solver version 2.7.0 (backward differentiation formula with absolute and relative error tolerances of 10^−10^) and the time series of biomass densities were saved for last 10,000 time steps. For numerical reasons, a local population was considered extinct and was set to 0 once its biomass density dropped below 10^−20^.

### Equations and parameters

For detailed equations and parameters, see Ryser et al. 2019^28^ and the Supplementary Material.

### Setup

To answer our questions, we model the following scenarios:

*Nutrient enrichment*: Simulations across a gradient of nutrient supply concentrations (0, 10) on one patch without emigration and therefore also no dispersal loss.
*Drainage effect*: Simulations across a gradient of maximal emigration rates (0, 0.15) on one eutrophic patch with a nutrient supply concentration of 10.
*Hostility effect with two patches*: Simulations across a gradient of dispersal losses (0, 1) on two eutrophic patches with a nutrient supply concentration of 15 on each and a maximal dispersal rate of 0.05.
*Heterogeneity effect with two patches*: Simulations across a gradient of nutrient supply concentrations (0, 15) on one of two patches with the other patch being a eutrophic patch with a nutrient supply concentration of 15, a maximal emigration rate of 0.05 and no dispersal loss.
*Interaction of hostility effect and heterogeneity effect*: For each level of heterogeneity (difference in nutrient supply between the two patches) we simulated the whole gradient of the hostility effect (dispersal loss of the predator from 0 to 1).
*Heterogeneity effect on complex food webs in complex landscapes*: For a complex meta-food-web, we generated 5 random geometric graphs consisting of 50 patches. Each patch was initialised with a complex food web consisting of 10 plant and 30 animal species. For all random geometric graphs, we simulated 15 homogeneous landscapes, where all patches have the same nutrient supply concentration with simulations across a gradient of nutrient supply concentrations ranging from 10^−0.8^ (oligotrophic) to 10^2^ (eutrophic) in steps of 0.2 in the exponent, and 5 heterogeneous landscapes, where the nutrient supply concentration for each patch is assigned randomly from the same gradient as in the homogeneous scenario.

