## Supplement for "Landscape heterogeneity buffers biodiversity of meta-food-webs under global change through rescue and drainage effects"

### Model

The model has been adapted from Ryser et al. 2019[1]. The feeding links (i.e. who eats whom) are constant over all patches and are as well as the feeding dynamics determined by the allometric food web model by Schneider et al. 2016[2] We integrate dispersal as species-specific biomass flow between habitat patches. Using ordinary differential equations to describe the feeding and dispersal dynamics, the rate of change in biomass density $B_{i,z}$ of species *i* on patch *z* is given by

$$\frac{dB_{i,z}}{dt} = B_{i,z} \sum_j e_j F_{ij,z} - \sum_j B_{j,z} F_{ji,z} - x_i B_{i,z} + E_{i,z} - I_{i,z} \quad \text{(for animals)} \quad (1)$$

$$\frac{dB_{i,z}}{dt} = r_i G_i B_{i,z} - \sum_j B_{j,z} F_{ji,z} - x_i B_{i,z} \quad \text{(for plants)} \quad (2)$$

with the first three terms describing local trophic dynamics and the last two terms describing emigration, $E_{i,z}$ (equation 9), and immigration, $I_{i,z}$ (equation 11). For simplicity, we do not let plants disperse. Trophic dynamics are driven by following three processes. First, predation or herbivory on species *j* with assimilation efficiency *e* ($e_j$ = 0.545, if *j* is a plant, typical for herbivory; $e_j$ = 0.906 if *j* is an animal, typical for carnivory[3]) and the functional response $F_{ij,z}$ (equation 3) for animals, and a nutrient dependent growth (equation 7) for plants. Second, losses due to predation or herbivory respectively. Third, losses by metabolic demands with $x_i = x_A m^{-0.305}$ with scaling constant $x_A = 0.141$ (tenfold laboratory metabolic rate[4] at a temperature of 20° Celsius to represent field metabolic rates) for animals and $x_i = x_P m^{-0.25}$ with $x_P = 0.138$ for plants. We used a dynamic nutrient model (equation 8) as the energetic basis of our food web. Each species *i* is fully characterized by its average adult body mass $m_i$. Body masses determine the interaction strengths of feeding links as well as the metabolic demands of species. Data from empirical feeding interactions are used to parametrize the functions that characterize the optimal prey body mass and the location and width of the feeding niche of a predator[2]. From each $m_i$ a unimodal attack kernel, called feeding efficiency $L_{ij}$ is constructed which determines the probability of consumer species *i* to attack and capture an encountered resource species *j*. We model $L_{ij}$ as an asymmetrical hump-shaped Ricker's function (equation 5) that is maximized for an energetically optimal resource body mass (optimal consumer-resource body mass ratio $R_{opt} = 100$) and has a width of $\gamma$. The maximum of the feeding efficiency $L_{ij}$ equals 1. Table TS1 is an overview of the standard parameter set for the equations. See also Schneider et al. 2016[2] for further information regarding the allometric food web model.

*Functional response*

$$F_{ij,z} = \frac{\omega_i b_{i,j} R_{j,z}^{1+q}}{1 + cA_{i,z} + \omega_i \sum_k b_{ik} h_{ik} R_{k,z}^{1+q}} \cdot \frac{1}{m_i} \qquad (3)$$

Per unit biomass feeding rate of consumer *i* as function of its own biomass density, $A_i$, (taking interference competition *c*, which is the time lost due to intraspecific encounters,), and biomass density of the resource $R_j$, with $b_{i,j}$, resource specific capture coefficient (equation 4); $h_{i,j}$, resource-specific handling time (equation 6); $\omega_i$ = 1/(number of resource species of *i*), relative consumption rate accounting for the fact that a consumer has to split its consumption if it has more than one resource species.

*Capture coefficient*

$$b_{ij} = f a_k m_i^{\beta_i} m_j^{\beta_j} L_{ij} \tag{4}$$

Resource specific capture coefficient of consumer species *i* on resource species *j* scaling the feeding kernel $L_{ij}$ by a power function of consumer and resource body mass, assuming that the encounter rate between consumer and resource scales with their respective movement speed. We differentiate between carnivorous and herbivorous interactions with each comprising a constant scaling factor for their capture coefficients $a_k$ with $k \in 0, 1$ ($a_0$ = 15 for carnivorous species and $a_1$ = 3500 for herbivorous species). For plant resources, $m^{\beta j}$ was replaced with the constant value of 1 (as plants do not move).

*Feeding efficiency*

$$L_{i,j} = \left(\frac{m_i}{m_j R_{opt}} e^{1-\frac{m_i}{m_j R_{opt}}}\right)^{\gamma} \tag{5}$$

The probability of consumer *i* to attack and capture an encountered resource *j* (which can be either plant or animal), described by an asymmetrical hump-shaped curve (Ricker's function), with width γ centered around an optimal consumer-resource body mass ratio $R_{opt}$ = 100 [2].

*Handling time*

$$h_{ij} = h_0 m_i^{\eta_i} m_j^{\eta_j} \quad (6)$$

The time consumer *i* needs to kill, ingest and digest resource species *j*, with scaling constant $h_0$ = 0.4 and allometric exponents $\eta_i$ = −0.48 and $\eta_j$ = −0.66.

*Growth factor for plants*

$$G_i = \frac{N}{K_i + N} \quad (7)$$

Species-specific growth factor of plants determined dynamically by the nutrient; with $K_i$, half-saturation densities determining the nutrient uptake efficiency assigned randomly for each plant species *i* and (uniform distribution within (0.1, 0.2)). The term in the minimum operator approaches 1 for high nutrient concentrations.

*Nutrient dynamics*

$$\frac{dN_z}{dt} = D(S - N) - \sum_{i,z} r_i G_i P_{i,z} \quad (8)$$

Rate of change of nutrient concentration $N$ of nutrient on patch $z$, with global turnover rate $D$ = 0.25, determining the rate at which nutrients are refreshed and the nutrient supply concentration $S$.

### Generating landscapes

We generated different fragmented landscapes, represented by random geometric graphs, by randomly drawing the locations of Z patches from a uniform distribution between 0 and 1 for x- and y-coordinates respectively.

### Dispersal

We model dispersal between local communities as a dynamic process of emigration and immigration, assuming dispersal to occur at the same timescale as the local population dynamics. Thus, biomass flows dynamically between local populations and the dispersal dynamics directly influence local population dynamics and vice versa. We model a hostile matrix between habitat patches that does not allow for feeding interactions to occur during dispersal. The total rate of emigration of animal species $i$ from patch $z$ is

$$E_{i,z} \ = d_{i,z} B_{i,z} \quad (9)$$

with $d_{i,z}$ as the corresponding per capita dispersal rate. We model $d_{i,z}$ as

$$d_{i,z} \frac{a}{1+e^{b(x_i - v_{i,z})}} \quad (10)$$

with $a$, the maximum dispersal rate, $b$ = 10, a parameter determining the shape of the dispersal rate, $x_i$, the inflection point determined by the metabolic demands per unit biomass of species $i$, and $\upsilon_{i,z}$, the per capita net growth rate of species $i$ on patch $z$. We chose to model $d_{i,z}$ as a function of each species' per capita net growth rate to account for emigration triggers such as resource availability, predation pressure and inter- and intraspecific competition. If for example an animal species' net growth is positive, there is no need for dispersal and emigration will be low. However, if the local environmental conditions deteriorate, the growing incentives to search for a better habitat increase the fraction of individuals emigrating.

**Immigration**

The rate of immigration of biomass density of species $i$ into patch $z$ follows

$$I_{i,z} = \sum_{n \in N_z} E_{i,n}(1 - \delta_{i,nz}) \frac{1 - \delta_{i,nz}}{\sum_{m \in N_n} 1 - \delta_{i,nm}} \qquad (11)$$

where $N_z$ and $N_n$ are the sets of all patches within the dispersal range of species $i$ on patches $z$ and $n$, respectively. In this equation, $E_{i,n}$ is the emigration rate of species $i$ from patch n, $(1 - \delta_{i,nz})$ is the fraction of successfully dispersing biomass, i.e. the fraction of biomass not lost to the matrix, and $\delta_{i,nz}$ is the distance between patches n and z relative to species i's maximum dispersal distance $\delta_i$ (see below paragraph Maximum dispersal distance). The term $\frac{1 - \delta_{i,nz}}{\sum_{m \in N_n} 1 - \delta_{i,nm}}$ determines the fraction of biomass of species $i$ emigrating from source patch n towards target patch $z$. This fraction depends on the relative distance between the patches, $\delta_{i,nz}$, and the relative distances to all other potential target patches m of species *i* on the source patch *n*, $\delta_{i,nm}$. Thus, the flow of biomass is greatest between patches with small distances.

For numerical reasons, we did not allow for dispersal flows with $I_{i,z} < 10^{-10}$. In this case, we immediately set $I_{i,z}$ to 0. We assume that the maximum dispersal distance $\delta_i$ of animal species increases with their body mass. For animal species, the body mass $m_i$ determines how far they can travel through the matrix. Thus, animal species at high trophic positions can disperse further than smaller animals at lower trophic levels. Each animal species perceives its own dispersal network dependent on its species-specific maximum dispersal distance

$$\delta_i = \delta_0 m_i^{\epsilon} \qquad (12)$$

where the exponent $\varepsilon = 0.05$ determines the slope of the body mass scaling of $\delta_i$. We chose a positive value for $\varepsilon$ to account for a higher mobility of animals with larger body masses.

### Table of parameters

**TS1**

| Symbol | Parameter | Value |
|---|---|---|
| $e_A$ | Conversion efficiency animal species | 0.906 |
| $e_P$ | Conversion efficiency plant species | 0.545 |
| $x_A$<br>exp | Scaling constant and exponent metabolic rate animal species | 0.141<br>-0.305 |
| $x_P$<br>exp | Scaling constant and exponent metabolic rate plant species | 0.138<br>-0.25 |
| $c$ | Interference competition | 0 |
| $a_0$ | Scaling factor capture coefficient for carnivorous links | 15 |
| $a_1$ | Scaling factor capture coefficient for herbivorous links | 3500 |
| $\beta_i$; $\beta_j$ | Allometric exponent for encounter rates | Carnivorous: 0.42; 0.42<br>Herbivorous: 0.19; 1 [5] |
| $R_{opt}$ | Optimal consumer-resource body mass ratio | 100 |
| $\gamma$ | Exponent Ricker's function | Foodchain: 2<br>Foodweb: 6 |
| $h_0$ | scaling factor handling time | 0.4 |
| $\eta_i$<br>$\eta_j$ | Allometric exponent handling time ($i$: consumer, $j$: resource) | -0.48<br>-0.66 [6] |
| $q$ | Hill coefficient | Foodchain: 1<br>Foodweb: 1.1 |
| $K$ | Half saturation density for nutrient uptake | Foodchain: 0.1<br>Foodweb: (0.1,0.2) |
| $D$ | Nutrient turnover rate | 0.25 |
| $S$ | Nutrient supply concentration | variable |
| $d_{max}$ | Maximum dispersal distance | 0.5 |
| $\varepsilon$<br>$D_0$ | Scaling factor and exponent for species-specific dispersal distance | 0.05<br>0.1256 |
| $a_S$ | Maximal emigration rate | Variable (Fig2b main text), 0.05 |
| $b$ | Shape parameter of emigration function | 10 |
| $f$ | Additional scaling factor for capture rates for stability | 0.05 |

### Results

#### *Rescue effect*

Increased dispersal loss (hostility) or the coupling with an oligotrophic patch (heterogeneity) essentially increases the strength of the drainage effect from the perspective of a eutrophic patch. However, while heterogeneity also increases the strength of the rescue effect from the perspective of an oligotrophic patch (Fig.S1, left to right), dispersal loss decreases the strength of the rescue effect (Fig.S1, bottom to top) except at high heterogeneity where the pattern is slightly more complex. Here (Fig.S1, top-left), the weakened coupling with a eutrophic patch induces oscillations (see section on dynamical interference).

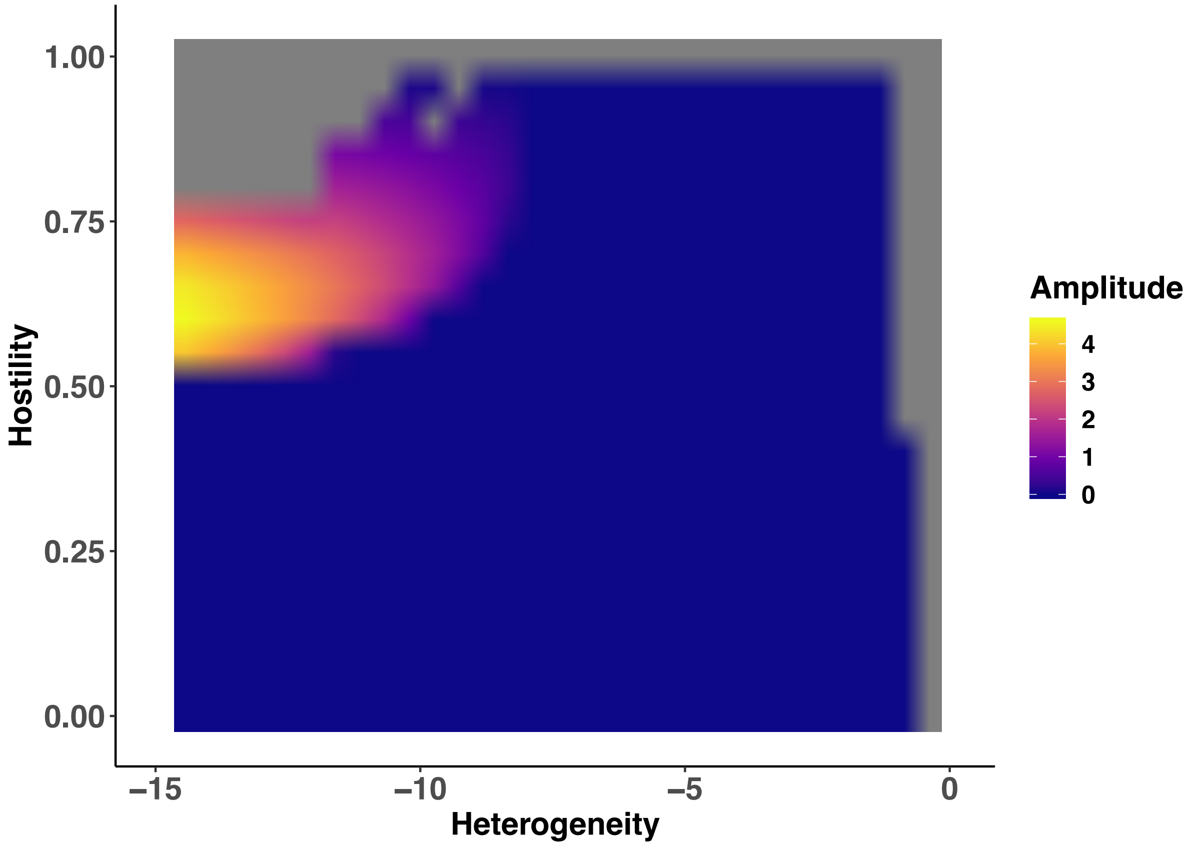


**Fig. S1:** Heat map showing the amplitude of biomass density oscillations in the predator (z-axis; colour coded) on the (always) oligotrophic patch across gradients of landscape heterogeneity (x-axis; difference in nutrient supply concentration between the two patches) and matrix hostility (y-axis). Amplitudes of 0 (blue) stand for an equilibrium state of the predator. Grey areas are where the predator went extinct.

*Dynamical interference*

When the hostility effect is very large, the coupling of the dynamics is weakened, which results in more chaotic oscillations as the frequencies get decoupled[7]. This in turn can lead to increased oscillations in the whole system that arise not from increased biomass fluxes but from dynamical interference (top quarter in Fig.3 and top-left corner in Fig.S1). This suggests that there is a lower threshold in strength of spatial links where instability arises from causes beyond the drainage and rescue effect. This becomes apparent in the top four rows in Fig.S2. As soon as the frequencies get decoupled, the reduction of amplitudes due do the drainage effect is overwritten and amplitudes increase again on the eutrophic patch (red).

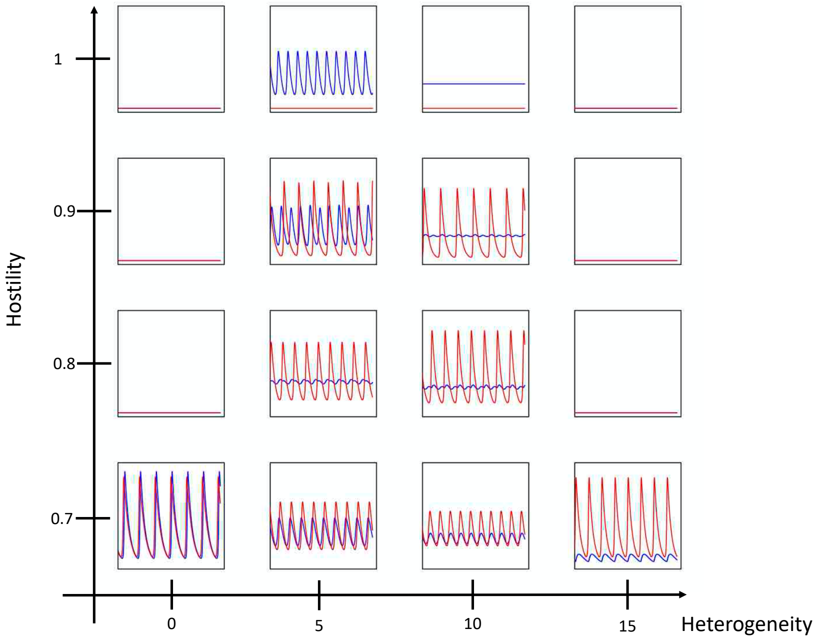

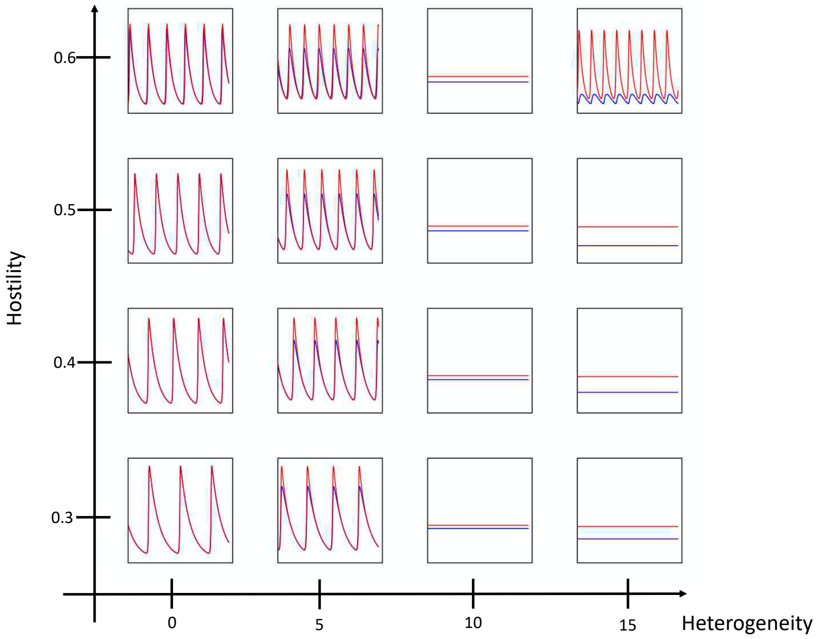


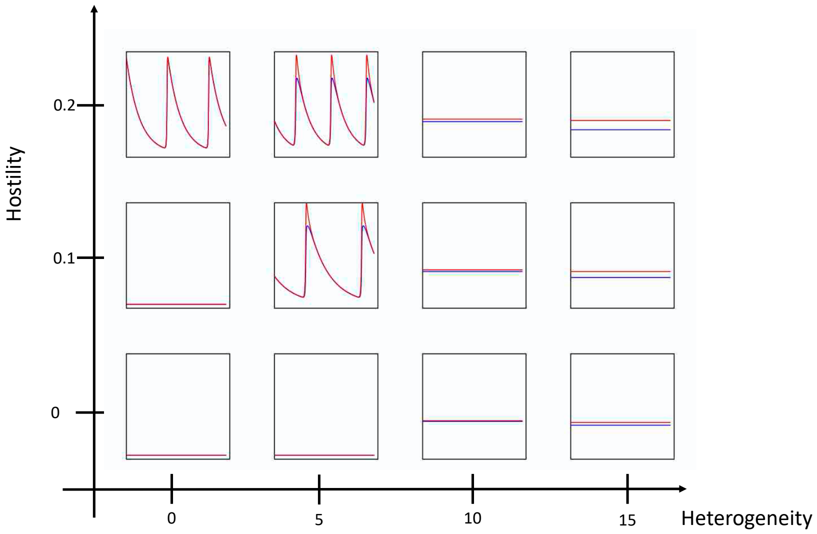


**Fig. S2**

Each plot represents biomass densities of the predator (y-axis) over time (x-axis) on the eutrophic patch (red) and on the variable patch (blue). Plots are arranged in a grid with the x-axis representing the landscape heterogeneity (delta nutrient supply of the eutrophic and the variable patch) and the y-axis representing the matrix hostility corresponding to Fig. 3 in the main manuscript.
